# The leading region of many plasmids is adapted for translational efficiency

**DOI:** 10.1101/2025.04.03.647016

**Authors:** Liam P. Shaw, Arancha Peñil Celis, Manuel Ares-Arroyo, Lijuan Luo, Tatiana Dimitriu, Fernando de la Cruz

## Abstract

To successfully transfer into a recipient, conjugative plasmids must overcome the new cell’s defense mechanisms. The leading region of many plasmids – the first part to enter the recipient – is highly enriched in anti-defense genes. Here, we investigate whether selective pressure from defense systems has also affected its sequence composition. We consider two possibilities: greater target depletion (to avoid triggering target-based defense systems) or greater codon adaptation (to more quickly express anti-defense genes). We analyse the leading region of 1,751 conjugative plasmids belonging to 13 plasmid taxonomic units (PTUs) present in the bacterial order Enterobacterales. First, we investigate GC content and possible depletion of short palindromic motifs (4-8bp) which could be a signature of selective pressure from Type II restriction-modification systems, and find evidence of depletion in certain PTUs. Then we use the codon adaptation index (CAI) to assess codon usage and find that different conjugative PTUs shown different patterns. We show that the leading region of plasmids with MOB-F and MOB-P relaxases is more adapted to optimal host codon usage, consistent with a selective pressure for translational efficiency in anti-defense genes that are transcribed while still single-stranded DNA. In particular, anti-restriction genes in the leading region have high CAI, and we use this fact to identify a single-stranded promoter in the F plasmid. Our findings uncover a new facet of plasmid evolution and emphasise the intensity of the selective pressure that plasmids face from defense systems.

## Introduction

To date, researchers have identified over 250 molecular systems which reduce or prevent infection by mobile genetic elements (MGEs) in bacteria (1). Whether these so-called ‘defense’ systems are conceptualised as a pan-immune system (2) or alternatively as weapons of inter-MGE conflict (3), any effective system will select for evasion in the MGEs it acts against. Most efforts on understanding evasion have focused on phage and their arms race with bacteria (4–8). Although it is established that plasmids carry anti-defense genes (9), far less is known about their strategies for evasion. Understanding plasmid evasion is important to understand horizontal evolution, as well as for synthetic biology approaches to engineer MGEs or to block the spread of plasmid-encoded antimicrobial resistance.

During conjugation, a plasmid is transferred from a donor into a recipient as single-stranded DNA (ssDNA) before circularisation and conversion to double-stranded DNA (dsDNA). This is a rapid process, taking just four minutes for the 100kb F plasmid (10), and always proceeds in the same way relative to the plasmid’s origin of transfer (*oriT*) (11). The leading region is the first portion of the plasmid to enter the recipient cell and is highly enriched in anti-defense genes (12). The immediate expression of these genes during transfer, identified in the 1990s and referred to as ‘zygotic induction’ (13), appears to be crucial for plasmid establishment (14).

Selective pressure from defense systems may also shape the leading region’s sequence as well as its gene content. Here, we consider two ways: greater target depletion (to avoid triggering target-based defense systems) or greater codon adaptation (to express anti-defense genes more quickly). To investigate both these possibilities in a dataset of well-characterised hosts, we tested them in 1,751 plasmids from within the bacterial order Enterobacterales, including those carrying multiple AMR genes of clinical relevance such as *bla*_NDM-1_, *bla*_CTX-M_ and *bla*_OXA-48_.

For target avoidance, we analysed the targets of the most common defense systems, Type II restriction-modification (RM), which are present in around 40% of bacterial genomes and mostly target 4-8bp palindromic motifs (1). In a previous study, we showed that plasmids tend to be depleted in Type II RM targets consistent with their known taxonomic range (15), and here we followed that approach to test whether this target depletion is stronger in the leading region.

For codon adaptation, our starting point is that speed matters during plasmid transfer. Previous work on the leading region has emphasised the rapid transcription of its anti-defense genes while still ssDNA (10,12,14,16,17), but the subsequent step of translation has been overlooked. This is despite these processes having similar rates: assuming maximal efficiency of ~85 nucleotides per second (18), a 1kb gene in *E. coli* may take 10s to be transcribed into mRNA and 10s to be translated into protein. This may seem short, but it is ~10% of the conjugative transfer time of the F plasmid, and inefficient translation will further delay it. Translation can be made more efficient through codon adaption: most amino acids can be specified by different codons, and certain codons lead to both faster elongation and greater mRNA stability (19). These optimal codons can be defined by examining a set of highly expressed chromosomal genes. We used this approach, quantifying codon usage with an established method (20) and comparing the region to the rest of the plasmid.

Our main finding is that the sequence of many leading region shows differences in codon usage compared to the rest of the plasmid, suggesting adaptation for translational efficiency.

Leading region genes have a higher codon adaptation index (CAI) and anti-restriction genes in particular have significantly higher scores. Using this improved understanding, we were able to identify a single-stranded promoter in the leading region of the F plasmid. Analysing codon usage may help identify novel features of plasmids, including new anti-defense genes, and help optimise novel conjugative delivery systems.

## Methods

We retrieved 19,017 bacterial plasmids from RefSeq200 and clustered them into related groups of Plasmid Taxonomic Units (PTUs) (21). We selected the fifty most prevalent PTUs consisting of 4,753 plasmids across 37 genera for initial analysis.

### Detection of leading regions

We followed a previously described procedure to identify leading regions (22). We first annotated all plasmids with prodigal v2.6.3 with the -p meta flag (23) and then used CONJScan v2.0 (24) to detect those with a relaxase, and thus able to be mobilised by conjugation. We discarded 16 PTUs that seemed incapable of moving by conjugation: 11 in which no conjugative trait was found (749 plasmids) and 5 phage-plasmids (279 plasmids). For the remaining 34 PTUs (3,725 plasmids) we screened for experimentally validated *oriT* sequences, detecting at least one *oriT* in 2,546 plasmids across 20 PTUs. To accurately identify the leading region of the plasmids, we discarded 336 plasmids which contained >1 *oriT* and/or >1 relaxase (except for PTU-X1 and X3, known to have two functional copies of the *oriT*_R6K_) (8), retaining 2,210 across 19 PTUs. Although there is no fixed definition for the size of the leading region, it is typically taken to be at least 10kb e.g. for the F plasmid it is ~13kb (9, Couturier). We therefore decided to only analyse plasmids >20kb, which led to discarding six small mobilizable PTUs from further analysis (median size 4.3kb, maximum 12.9kb). We also discarded PTU-HI2 since >50% of the plasmids within it lacked a known *oriT*. In summary, we could confidently identify a leading region in 1,751 conjugative plasmids across 13 PTUs, the vast majority of which (1,736, 99.1%) were from within Enterobacterales (Table S1).

### RM system prevalence

For each of the 37 genera with plasmids in our initial dataset, we downloaded a random sample of at most 100 complete bacterial genomes from RefSeq (3,335 genomes) and then used rmsFinder v1.0.0 (https://github.com/liampshaw/rmsFinder) with default thresholds to predict the presence of Type II RM systems. We detected 1,335 systems with confidently predicted motifs in 1,020 genomes, with 77 unique targeted motifs corresponding to 213 motifs after deambiguation (e.g. CCWGG = {CCTGG, CCAGG}). The majority of RM motifs were of length k=5 and k=6 (64/76, 83%); within Enterobacterales, there were 89 such motifs across 17 genera, with a median prevalence of 2% per genera. We used this to produce a database of RM systems with known targets across genera (Table S2).

### Depletion of RM motifs

Palindromes have previously been used as a proxy for Type II RM targets (6,7,15), and we also use this approach here for initial analysis. Because a DNA palindrome of length k is determined by each of its three bases matching with its complement at the right position in the last three bases, the expected density of palindromic k-mers in a long sequence with a GC content of *p* is given by:

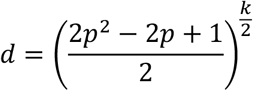

For example, for a sequence with GC content of 50% we get d=(1/4)^3^=0.01565 per basepair, corresponding to a palindrome density per kilobase of ~15.6. We use a sliding window over plasmids to compare the observed number of total palindromes to this expectation.

We also used RMES v3.1.0 (25) to calculate a more rbust representation score for all *k*-mers (sequence motifs of length *k*). RMES (https://forgemia.inra.fr/sophie.schbath/rmes) compares the expected number of occurrences of a *k*-mer to the observed occurrences to calculate a representation score, with negative scores indicating under-representation. We used the maximal model which uses the (*k-*1)-mer sequence composition to calculate expected occurrences, constituting the most conservative version because it uses as much information about sequence composition as possible. We compared the leading and lagging regions of plasmids with a fixed length of 10kb to control for sequence length.

### Gene prediction and functional annotation

After finding protein-coding genes on all plasmids with prodigal v2.6.3 with the -p meta flag (23) we performed functional annotation with eggNOG-mapper (version emapper-2.1.13) (26) based on eggNOG orthology data (27) and diamond for sequence searches (28). 79.7% of predicted proteins had a hit (163,961 of 205,673) and 63.3% (130,171) were assigned to an informative COG one-letter code (i.e. excluding S, R and −). We detected AMR with AMRFinderPlus (software v4.2.7, db v2026-03-24.1) with default parameters (>0.5 coverage, >0.9 identity) (29). We searched for defense and anti-defense systems with DefenseFinder v2.0.1 (1,9) using defense-finder-models v2.0.2 and CasFinder v3.1.0. This identified 1,858 defense genes and 4,397 anti-defense genes in 1,617 and 3,174 systems respectively (Table S3 gives 7,004 individual HMM hits with evalue<10^−10^). We searched for tRNAs in all plasmids with trnascan-SE v2.0.12 (30) to confirm that plasmids were dependent on host tRNAs for expression. Only three plasmids had non-pseudo tRNAs reported: NZ_CP017387.1 (PTU-C), NZ_CP030077.1 (PTU-C), and NZ_CP040265.1 (PTU-FE). These are likely the result of previous chromosomal integration and excision.

### Codon usage analysis

The optimal set of codons for an organism are those that occur most frequently in genes translated at high abundance for a given amino acid i.e. for a given amino acid, there is a optimal codon. These optimal codons get a weight *w*=1, and synonymous codons for the same amino acid get a weight defined by their relative usage frequency to the optimal codon. The Codon Adaptation Index (CAI) of a gene is defined as the geometric mean of the w-values of its *M* codons (20):

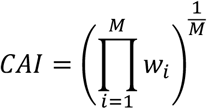

CAI ranges from 0 to 1, with 1 corresponding to optimal codon usage. Since n=892 (50.9%) plasmids were from *Escherichia*, we used the *E. coli* codon weights table calculated by Lucks et al. (31) based on 27 very highly expressed genes including the ribosomal genes (32). For context, the median CAI in the *E. coli* K12 genome is 0.321, and values range from 0.066 (*mgtL*) to 0.830 (*lpp*, from the set of genes used to define optimal codons). We verified that we could recalculate the 61 codon weights from *E. coli* K12 MG1655 matching the figures from Lucks et al. (31), then also calculated the analogous weights for other species using blast hits in reference genomes for the 27 highly expressed genes (Table S4). Within Enterobacterales we found high correlation with the *E. coli* codon weights: *Salmonella enterica* (GCF_000006945.2) Pearson’s r=0.987, *Klebsiella pneumoniae* MGH78578 (GCF_000016305.1) r=0.986, *Citrobacter freundii* (GCF_904859905.1) r=0.997, *Enterobacter hormaechei* (GCF_019048245.1) r=0.947. We calculated CAI for all ORFs found with prodigal in plasmids (Table S5).

## Results

### Leading regions are associated with anti-defense genes

We defined the leading region as the sequence following the *oriT* in opposite direction to the relaxase i.e. the start of the T-strand (Fig. 1a, see Methods). We could confidently locate the leading region in 13 conjugative PTUs within Enterobacterales (1,751 plasmids of which 1,736 from Enterobacterales, median length 91kb range 21.9-329.3kb, median length per PTU 42-217kb). Conjugative transfer of ssDNA induces defense mechanisms such as the bacterial SOS response (33), but many plasmids can carry anti-defense genes that are expressed early in conjugation. These include *ardA*, encoding an anti-restriction protein that inhibits the endonuclease activity of Type I systems (34), or *psiB*, encoding a protein which inhibits the SOS response (35). As previously reported, the leading region is enriched in these anti-defense genes (12) and we reproducde this result in our dataset (Fig. 1b). We noted that anti-CRISPR systems were further from the *oriT*, suggesting CRISPR may take longer to act once the plasmid enters. We hypothesised that selective pressure from defense systems could also lead to distinct sequence composition as well as gene content (Fig. 1c). It should be noted that anti-defense enrichment patterns differ by relaxase type (Fig. 1d-f). PTUs where plasmids carried a MOB-F relaxase showed a consistent leading region peak in anti-defense systems, as did MOB-P relaxase PTUs, but MOB-H PTUs showed no detectable peak and some PTUs had no anti-defense systems detected. We return to these observations below.

**Figure 1.**
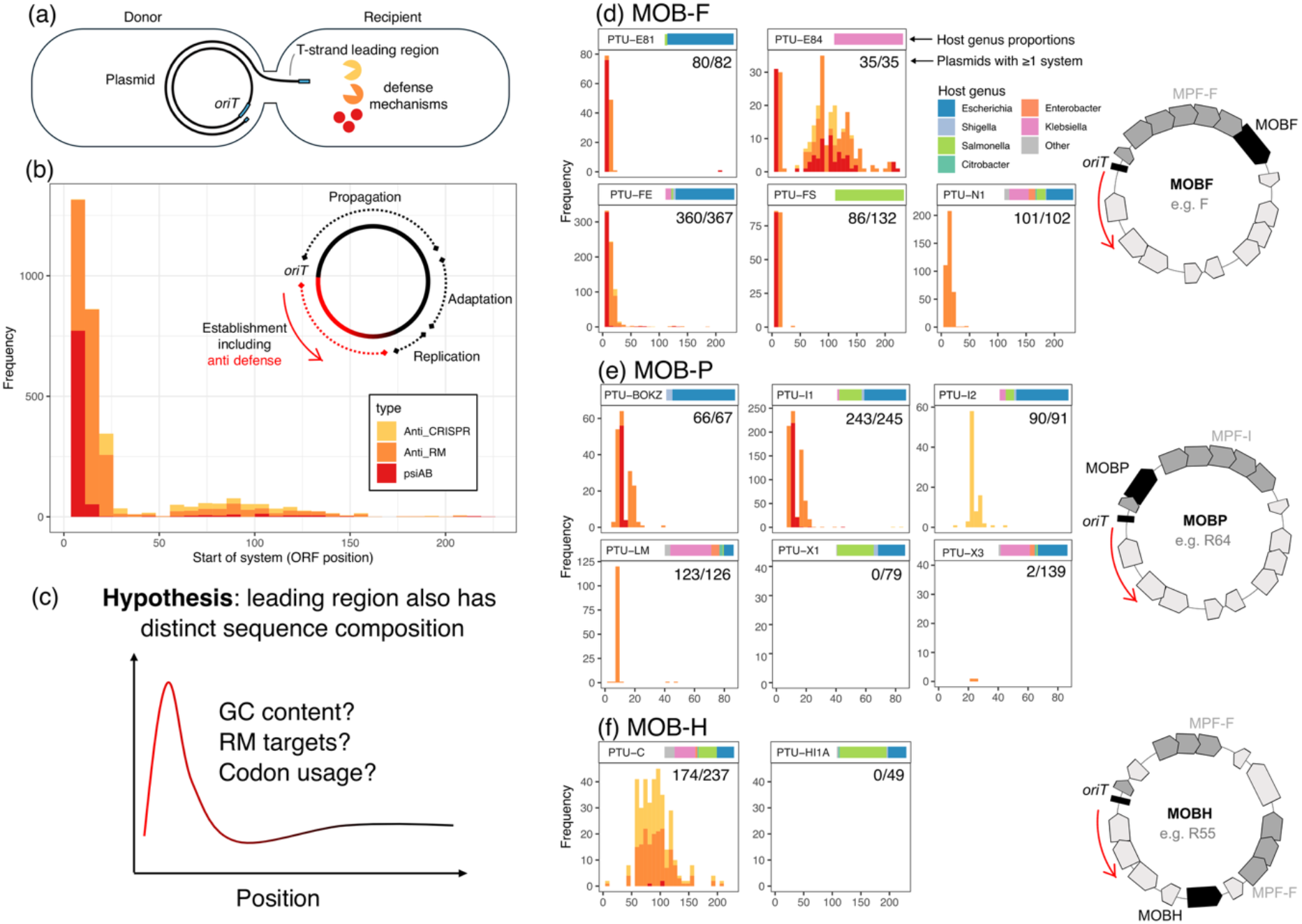
The leading region shows distinct gene content because of selective pressure from defense systems. (a) Schematic showing conjugative transfer of a plasmid from a donor into a recipient. The leading region on the transfer strand (T-strand) is under strong selective pressure from defense systems including RM, CRISPR-Cas and abortive infection. (b) The leading region is enriched in anti-defense genes, shown here for over 1,751 plasmids in 13 plasmid taxonomic units (PTUs) found mainly in Enterobacterales. Inset shows a schematic of plasmid organisation following Samuel et al. (12). (c) We hypothesise that this selective pressure from defense systems also shapes the leading region’s sequence composition. (d-f) Enrichment of anti-defense systems in the leading region (frequency i.e. absolute numbers), subsetted by mobility type (the most common relaxase detected within the PTU, with diagrams showing indicative plasmid organisation) and then by PTU. X-axis shows ORF position as in (b). Inset bars in facet titles show host genera proportions, numbers in top-right corner of plots give the number of plasmids with at least one anti-defense system detected. Some PTUs have no anti-defense systems detected (PTU-X1, PTU-HI1A). For individual anti-defense HMM hits (evalue<10^−10^) see Table S3. Not shown in figures (b) and e) is a single anti-Pycsar system detected in one PTU-BOKZ plasmid (NZ_CP023144.1).

### Evidence for GC richness and motif depletion

The leading region had higher mean GC content in many plasmids, with differences between PTUs (Fig. 2a-c). 10/11 PTUs associated with MOB-F and MOB-P relaxases had higher GC in the leading 25kb compared to the rest of the plasmid (Fig. 2d). GC skew in the transfer strand – which expresses the ratio between G and C on a single strand – also often had higher variance in the leading region (Fig. S1). Next, we investigated the possible impact of RM systems on leading region sequence composition. Because Type II RM targets tend to be short palindromes 4-8bp in length, we initially looked at the density of short palindromes across plasmids. We compared the palindrome density expected based on GC content with the observed density (see Methods). 4-bp and 8-bp palindromes showed no clear difference in the leading region (Fig. S2 and S3). Patterns of depletion in bacterial genomes are often strongest for 6-bp palindromes (36) as also previously established for plasmids (15), and interestingly the leading region had fewer 6-bp palindromes than expected in some PTUs, such as PTU-E84, PTU-FE, PTU-BOKZ and PTU-I1 (Fig. 3). However, many PTUs showed no signatures of depletion.

**Figure 2.**
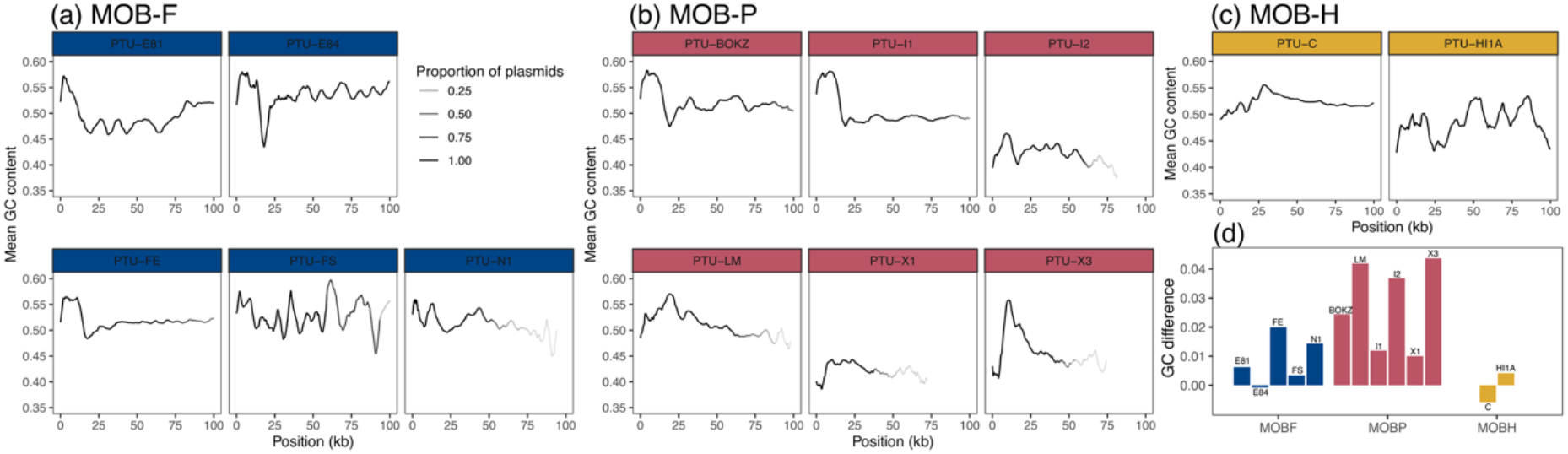
The leading region has higher GC content in many PTUs. Mean GC content in non-overlapping 1kb windows shown for (a) MOB-F (b) MOB-P and (c) MOB-H plasmids, faceted by PTU. Plots are truncated at 100kb with line intensity showing the proportion of plasmids being averaged over within the PTU at each position. (d) The difference in mean GC content between the leading 25kb and the rest of the plasmid for each PTU.

**Figure 3.**
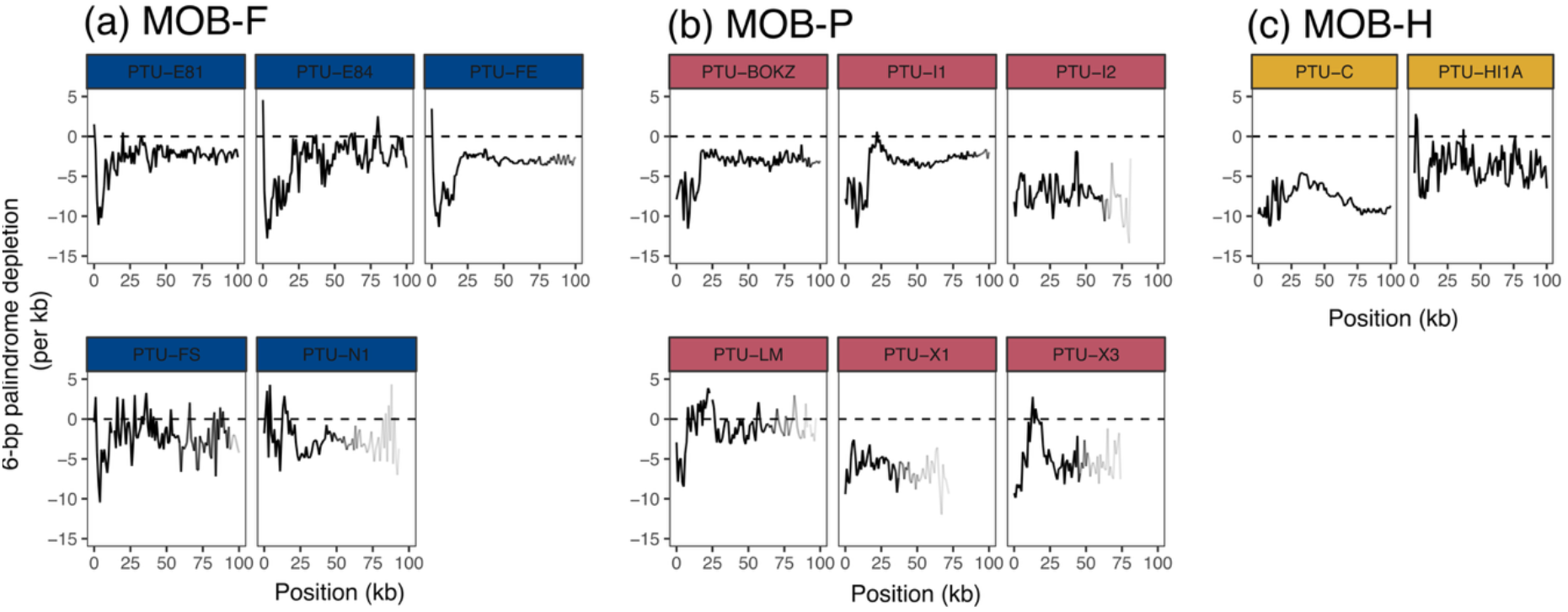
The leading region of some PTUs is depleted in 6-bp palindromes. The difference between observed palindrome density in nonoverlapping 1kb windows and the theoretical density expected from GC content (see Methods), such that lower values show depletion below expectation. Line intensity indicates proportion of plasmids being averaged over.

Averaging over plasmids and palindromes in this way might obscure real biological signal, since different PTUs have different hosts and are therefore subject to different selective pressures. We therefore performed an analysis for each PTU for the motifs targeted by RM systems within its observed species range (‘within-range targets’), using RMES to compare the number of observed motifs to the number expected under a null model that controls for sequence composition; negative/positive RMES scores correspond under/over-representation. Since only PTU-C had a within-range target of k=4, we only analysed targets of k=5 and k=6, comparing the leading and lagging region (Fig. 4). On average, the leading region was not depleted in within-range targets for k=5 but was for k=6 (scores <0 in 10/13 PTUs, one-tailed binomial test p=0.046), in line with previous work suggesting depletion applies predominantly to 6-bp motifs (12, 17).

**Figure 4.**
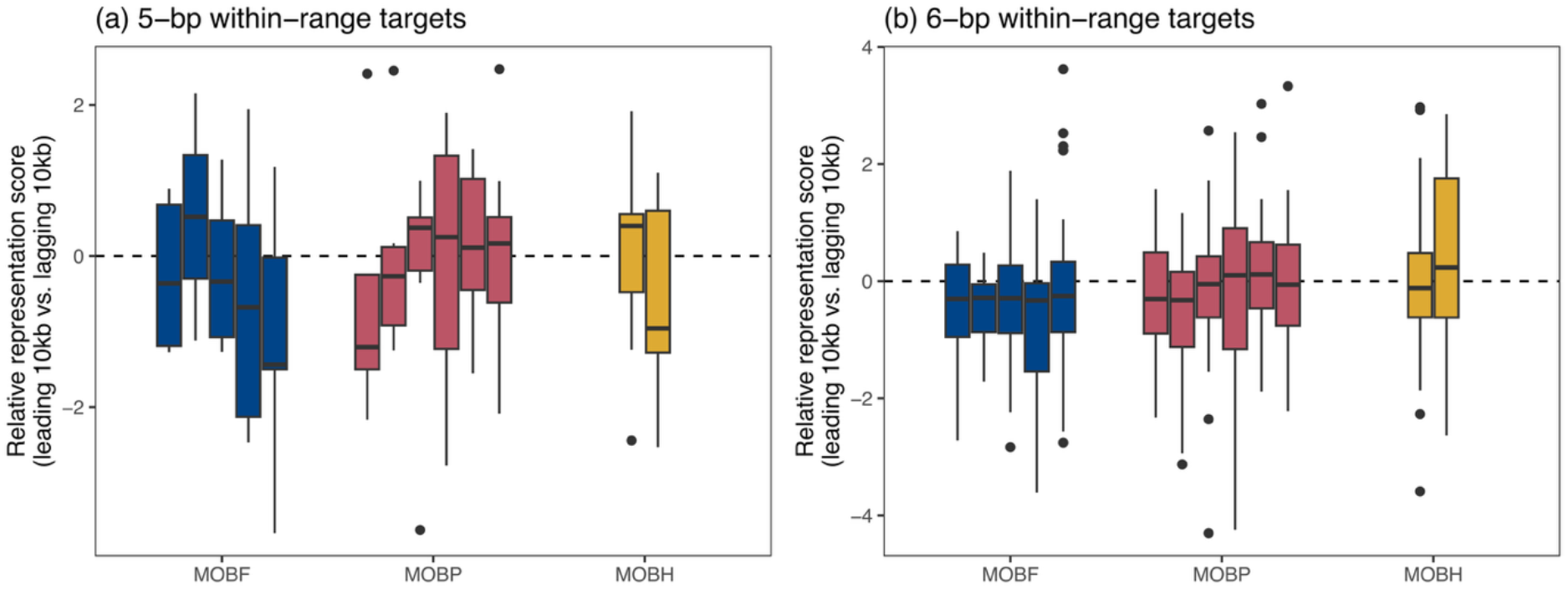
Relative representation scores for within-range RM targets in the leading vs. lagging regions of plasmids. Panels show mean results for (a) 5-bp and (b) 6-bp targets of mean differences in RMES score for the leading vs. lagging region (10kb each) of the within-range target across all plasmids within each PTU.

However, there is no significant mean depletion in the leading vs. lagging region across all within-range RM targets and targets (Fig. S4; mean depletion −0.14, Wilcox signed rank test p=0.08). Furthermore, these representation scores correspond to tiny changes in numbers of motifs. To take a specific example, consider PTU-B/O/K/Z. These plasmids are narrow-host-range, seen only in *Escherichia* and *Shigella*. Our analysis suggested that ~3% of *Escherichia* genomes contain Type II restriction enzymes similar to enzyme Cfr10I which target RCCGGY = (ACCGGC, GCCGGC, GCCGGT, ACCGGT), and indeed of these four motifs, the first three are the most depleted known targets in the leading region. However, in general we found that closer examination did not reveal strong specific signals, making us cautious about over-interpreting this weak average signature. Within Enterobacterales (n=17 genera), for targeted motifs of k=5 (32 motifs) and k=6 (89 motifs) the median motif was targeted by RM systems with only 2% prevalence (IQR 1-4%). The most prevalent case was in *Erwinia*, where 70% of genomes had a Type II RM system targeting GTATCC, but the only plasmid in our dataset from *Erwinia* (NC_005246.1) had 4 instances of this motif in its leading region. We therefore conclude that there is no evidence for systematic depletion of RM targets in the leading region.

As well as potential vulnerability from Type II RM systems, we also considered the possibility that palindromes in the leading region might form transient dsDNA hairpins that could also potentially be targeted by the nuclease SbcCD (37) as suggested by Fraikin et al. (11). We searched for near-perfect palindromic hairpins at least 4bp in length where the inverted repeats were separated by no more than 1kb (einverted in EMBOSS v6.6.0.0, flags: -threshold 10 -maxrepeat 1000 -mismatch -99 -gap 12 -match 3) but found no evidence for association of palindrome densities and position in the leading 50kb (Fig. S5).

### The leading region shows adapted codon usage

For each plasmid gene, we computed the Codon Adaptation Index (CAI) relative to optimal codon usage in *E. coli* (see Methods). We found a strong signal of differential sequence composition in the leading region, with higher CAI for positions in the leading region of PTUs with MOB-F and MOB-P relaxases (Fig. 5). When we tested across plasmids, we found that their anti-defense genes had higher CAIs than their other genes (Fig. 6a; median 0.34 vs. 0.27 for all other genes, Wilcoxon rank sum test p<0.001) and were even slightly above the CAI for the average *E. coli* chromosomal gene (median 0.34 vs. 0.32, p<0.001). Within the category of anti-defense, anti-restriction genes had significantly higher CAIs than anti-CRISPR (median 0.385 vs. 0.331) or *psiAB* (median 0.336, range 0.256-0.407). Analysing genes using the KEGG ontology (Fig. 6b) showed that genes with high CAIs made biological sense, including those encoding proteins for pilus formation, heavy metal resistance, iron binding and phage shock response, with anti-defense genes having higher than average CAI. In particular, anti-restriction genes encoding proteins in the ArdB/KlcA/AcrIC11 family, which were seen across 10 PTUs in 1,054 plasmids (60.7%), were highly optimised for codon usage (CAI 0.442, range 0.28-0.52). Furthermore, there was a significant negative correlation between distance from *oriT* on the leading strand and CAI for anti-defense genes (Pearson’s R=−0.259, p<0.001) which was much stronger than for other genes (Pearson’s R=−0.054, p<0.001).

**Figure 5.**
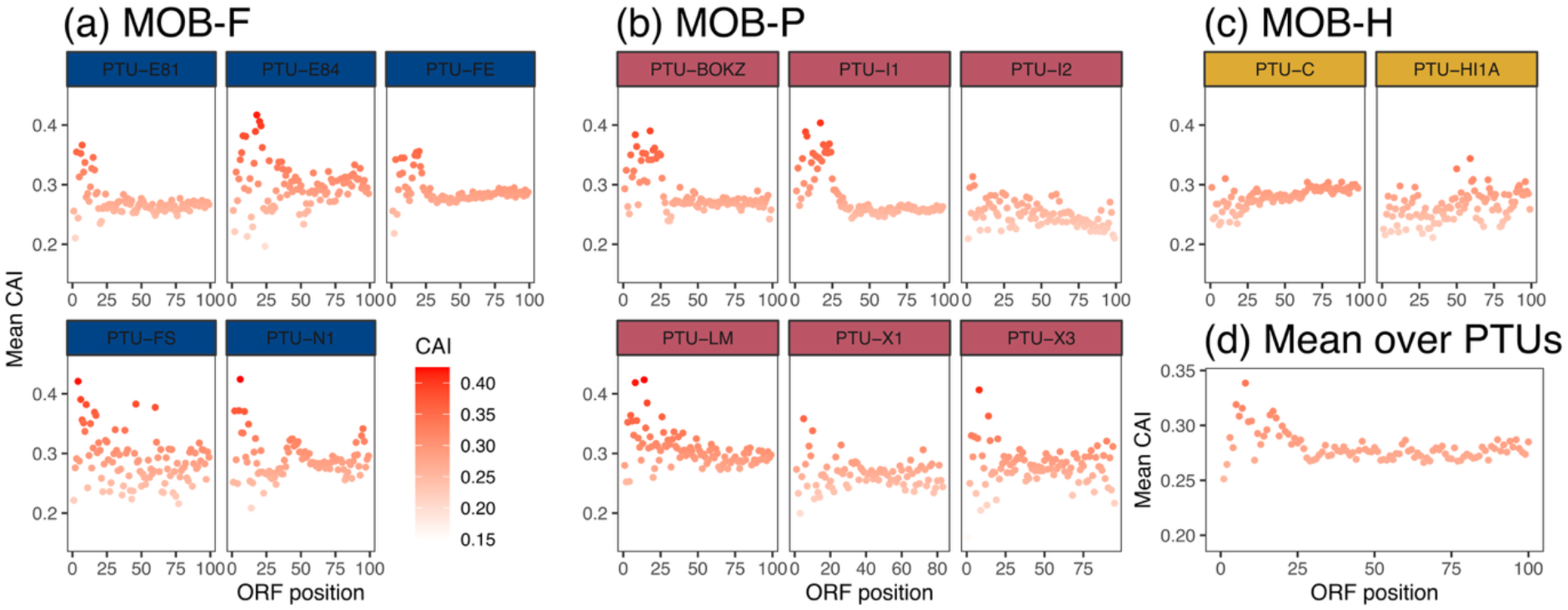
The leading region has a peak in host codon adaptation. (a)-(c) Plots of mean codon adaptation index (CAI) by ORF position for each PTU, with colour indicating mean CAI of genes found at that position. (d) Mean CAI for each position averaged over PTUs (i.e. each PTU given equal weighting). Plot truncated at 100 ORFs. CAI was calculated against *E. coli* codon weights determined from highly expressed genes (see Methods).

**Figure 6.**
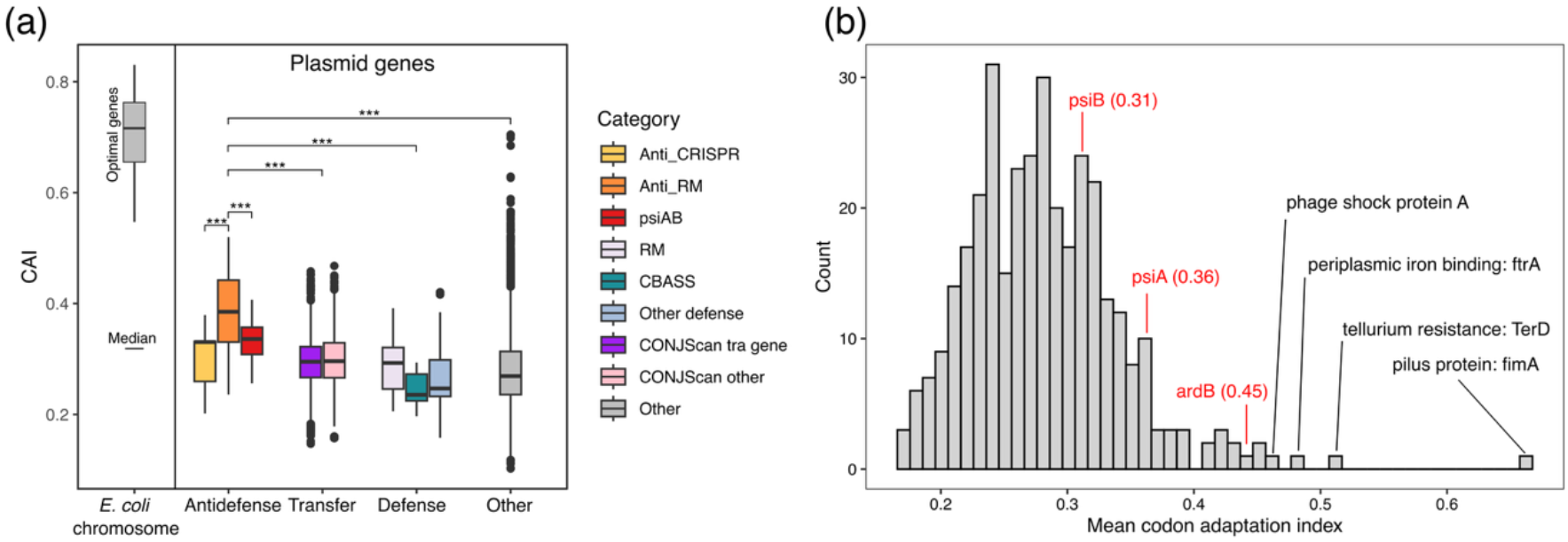
Anti-defense genes have more optimal codon usage than many other plasmid genes. (a) Histogram of plasmid genes from all 1,751 plasmids, after classifying into defense and anti-defense (DefenseFinder) and conjugative genes (CONJScan). Data for *E. coli* chromosomal genes is shown for comparison (27 optimal set of genes used to define optimal codon weights and median). Statistical tests are Wilcoxon rank sum tests, all p<0.001. (b) Histogram of mean CAI for KEGG KO categories with 10 or more gene representatives. Anti-defense genes like those encoding ArdB or PsiA/B are above average in the distribution, although there are some categories with more optimal codon usage.

### Codon usage patterns can help identify promoters

Historically the leading region of the F plasmid has been defined as ~13kb in size (38,39). We therefore analysed the F plasmid in detail to understand the patterns of CAI as a detailed case study (Fig. 7a). Genes on the transfer strand had higher CAI in the leading region (Fig. 7b), notably higher than the *tra* genes in the transfer region (Fig. 7c). The F plasmid has a known single-stranded promoter that allows transcription of genes in the leading region while they are still ssDNA. Couturier et al. used sequence similarity to the F*rpo* region (renamed F*rpo1*) to identify a single-stranded promoter (F*rpo2*) that shares 92% identity with F*rpo1*. (This region was previously noted by Masai and Arai in their original publication identifying F*rpo* (40).)

**Figure 7.**
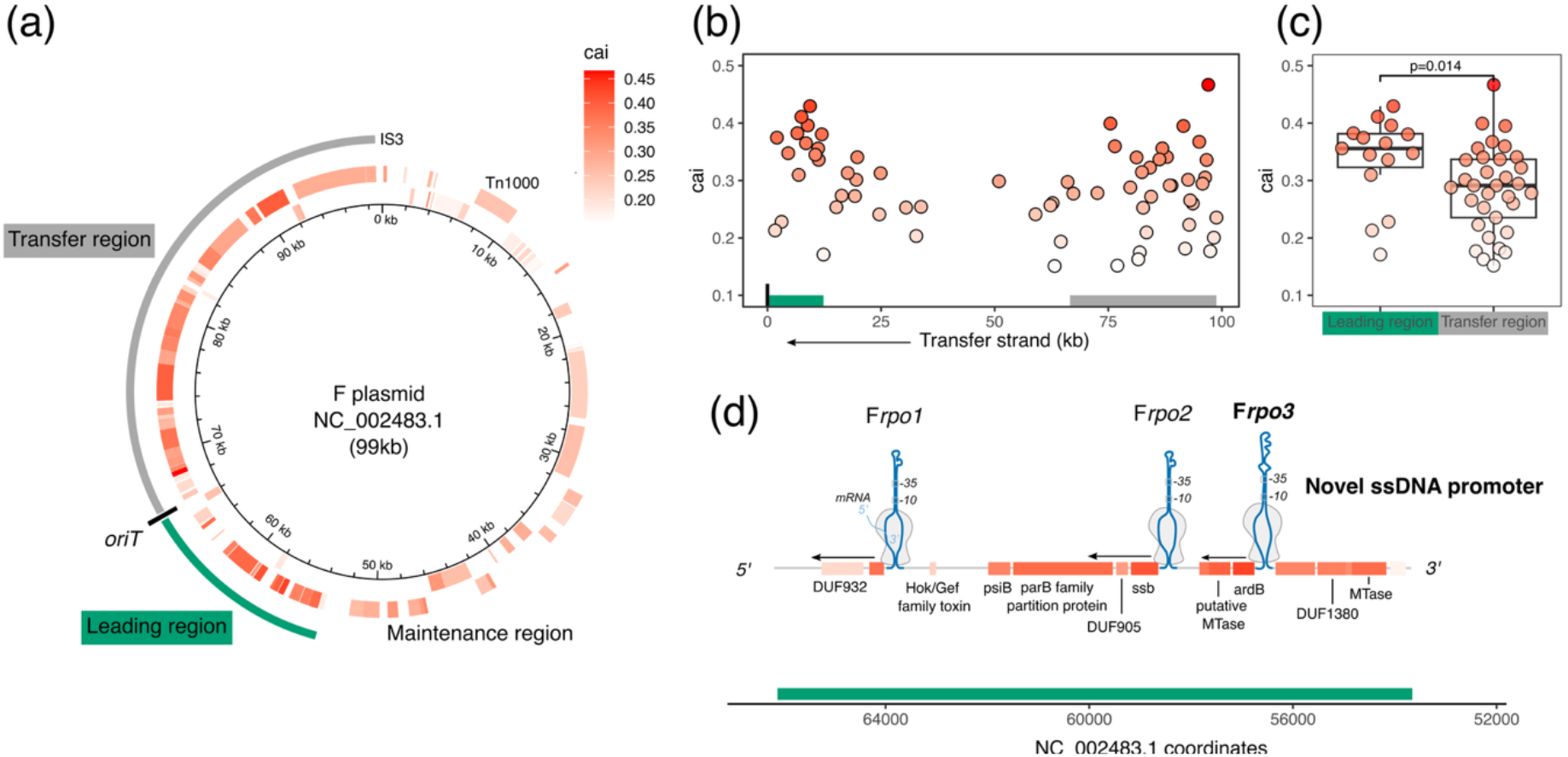
The leading region of the F plasmid is adapted for rapid expression. (a) F plasmid map based on NC_002483.1 using NCBI annotations (non-pseudogenes), with genes coloured by Codon Adaptation Index (CAI). (b) Visualisation of CAI relative to *oriT*, with sliding average of 15 ORFs shown in black (midpoint). Only genes encoded on the transfer strand are shown i.e. in the correct orientation for transcription upon entry into the recipient cell. (c) Leading region genes on the transfer strand have significantly higher average CAI than transfer region regions (Wilcoxon signed-rank test, p<0.05). (d) Single-stranded promoters in the leading region allow transcription of codon-adapted genes while still ssDNA. These include the known ssDNA promoters F*rpo1* and F*rpo2* and a ssDNA promoter which we name F*rpo3*, identified by inspecting the secondary structure upstream of the cluster of codon-adapted genes including anti-restriction gene *ardB* and a putative methyltransferase. Secondary structures for Frpo1/2 have been adapted from Figure 4 of Couturier et al. (10) under a Creative Commons License https://creativecommons.org/licenses/by/4.0/.

As expected, genes associated with these promoters have high CAI. However, examining the leading region in detail, we noticed that the cluster of genes including *ardB* and a putative methyltransferase had high CAI but no known promoter. Blast recovered no hits to F*rpo1* or F*rpo2*. However, when we computed the secondary structure for the DNA upstream of *ardB* on the transfer strand with mFold v3.6 (41) we were able to identify a putative single-stranded promoter, which we named F*rpo3*. (We also searched for the region upstream of the next cluster of genes along, but failed to identify a probable promoter.) The sequence of F*rpo3* is distinct from F*rpo1* and F*rpo2* but it forms a similar secondary structure: a long hairpin within we which we could identify a −10 and −35 box, making it plausible that it directs transcription along the transfer strand (Fig. S6). F*rpo3*, we found that it also shares 86% sequence identity with a putative ssDNA promoter identified by Samuel et al. from an insect gut metagenome (12). After initial publication of this paper as a preprint, we were made aware of recent work by Wen et al. using RNA-seq (42), which independently identified the region corresponding to F*rpo3* (‘class 2’ in their terminology). Further work by Fraiken et al. (43) shows that it also acts a ‘roadblock’ to dsDNA conversion which allows time for these early genes to be transcribed, providing evidence that plasmids tune gene expression and complementary strand synthesis in a unified strategy.

## DISCUSSION

The leading region of plasmids has been recently shown to be enriched in anti-defense genes. Here we show that it is also adapted to optimal codon usage. Our findings are consistent with a selective pressure for the high expression of anti-defense genes for plasmid establishment, and emphasise the importance of considering how evolution has shape plasmid genome organization.

We considered several possible adaptations in sequence composition. We found that the leading region of conjugative plasmids with MOB-F and MOB-P relaxases is GC-rich, with evidence of higher variance in GC skew on the transfer strand. To our knowledge this observation has not been discussed extensively in the literature, although one previous hypothesis has been made about structural requirements for DNA loading into the transfer channel using G-quadruplexes (22). We also found that there was an average depletion of 6-bp palindromes of ~5 per kilobase in the leading region across all plasmids, although this signal did not apply to all PTUs and was due to those with MOB-F relaxases like PTU-FE, PTU-BOKZ, and PTU-I1, which include many typical *E. coli* plasmids. We attempted to analyse the motifs targeted by RM systems that plasmids encounter most often. With the caveat that it is challenging to know how to accurately average over motifs targeted by diverse RM systems with very different prevalences and (unknown) efficiencies – or indeed what such an average really means – we found only a weak average signal of 6-bp depletion. This is consistent with previous work highlighting that depletion is largely a feature of smaller plasmids (<10kb) since larger plasmids cannot easily lose motifs by mutation, and seem to adapt instead by carrying additional genes for evasion (15).

Any depletion of RM targets might be somewhat surprising since the leading region enters the recipient as ssDNA and Type II RM systems generally target only dsDNA. While some Type II restriction enzymes can have activity against secondary structures formed by ssDNA (44) we found no general depletion of very short putative hairpin structures. Therefore, although it remains possible that individual instances of plasmid infection could be thwarted by motifs in the leading region, akin to a recent report for a dsDNA phage (45), overall our results lead us to conclude that there is no general pattern of Type II RM motif depletion in the leading region. The pattern of GC-richness and palindrome depletion in plasmids with MOB-F relaxases suggests different selective pressures to the rest of the plasmid, but the reasons are unclear. Since the leading region enters the recipient as ssDNA its secondary structure during this initial phase is probably crucial for its function. In general, GC-rich DNA forms more stable transient structures because of the higher thermodynamic stability of GC bonds, and future work might elucidate the possible effects on the leading region’s structure. It is interesting that the single-stranded promoter regions of the F plasmid are GC-rich (mean GC content of F plasmid is 48.2% but F*rpo1*: 58.0%, F*rpo2*: 56.7%, Frpo3: 68.6%). It seems probable that other aspects of secondary structure may be important: for example, GC content may be low at the very start of the transfer strand to reduce transient structures that impede transfer, before higher GC in the subsequent leading region.

Considering codon usage, we found that genes in the leading region of most PTUs were more adapted for translational efficiency than the rest of the plasmid, as assessed by the codon adaptation index (CAI). We stress that CAI is not a measure of a gene’s importance and that gene expression can be optimised in many other ways; in general, there is not a strong relationship between codon bias and translation efficiency because of the many other factors affecting both (19), and alternative measures of codon adaptation that account for mutation bias can also be used (46). Our interpretation of codon adaptation might apply to genes with a known dose-dependent response – for example, plasmid-borne beta-lactamases have higher CAI than other AMR genes (median 0.30 vs. 0.24, Wilcox signed rank test p<0.001).

However, speed is probably the most important factor. Genes controlled by ssDNA promoters with high CAI will translate quickly, helping with the rapid conversion to dsDNA, which is important to avoid increased induction of the SOS response. Speed would be expected to be most important for anti-restriction compared to other anti-defense genes, because preventing early restriction can allow subsequent methylation to occur, protecting the plasmid from further restriction. Indeed, the high CAI of the cluster of genes containing the broad anti-restriction gene *ardB* on the F plasmid’s transfer strand led us to identify a putative promoter, F*rpo3*, highlighting the value of considering the dynamics of zygotic induction. After publication of our results, we discovered that independent research had also identified this promoter (along with others in the leading region) and experimentally proved its function (42,43). Future work into the relationship of initiation and elongation along the transfer strand will lead to a better understand of how the leading region is organized in the F plasmid, and potentially beyond.

Although we used *E. coli* optimal host codon usage, fast-growing bacteria and highly-expressed genes are known to converge on a small subset of optimal codons and anticodons (47). Indeed, we found codon weights for other Enterobacterales species are highly correlated (r>0.9) so our analysis is not dependent on this assumption. Codon usage bias is also known to be stronger for organisms with multiple habitats (48). By analogy, the leading region of Enterobacterales-specific PTUs may represent a sequence optimised to deal with their multiple potential hosts. Previous work on codon usage in plasmids has largely considered the effect on host fitness, such as experiments in *E. coli* which found that altering host translational efficiency did not change plasmid costs (49), or effects on plasmid retention due to codon usage mismatch (50). By approaching the problem from the perspective of plasmid establishment, we argue that plasmid success depends not on general adaptation for translational efficiency, but specifically in the leading region.

Our results are consistent with literature that suggests that rates of horizontal gene transfer are correlated with the similarity of tRNA pools (51) and require codon usage compatibility (52). Our dataset also had n=15 PTU-C representatives found in species outside Enterobacterales and we note that codon weights for these species are not as strongly correlated with *E. coli* e.g. *Shewanella algae* (r=0.814), *Vibrio cholerae* (Pearson’s r=0.721) amd *Photobacterium damselae* (r=0.590) (based on 22 highly expressed genes in each reference genome). The lack of a peak in CAI in the leading region of PTU-C plasmids could be related to their broad host range, a sign of a potential tradeoff between specialisation and generalism, but this reasoning does not apply to the narrow host-range PTU-HI1A.

General statements about plasmids are based on two types of evidence – both have issues. On the one hand, bioinformatic analysis of many plasmids is used to make statements about the ‘average’ plasmid, often without acknowledging that this is usually just the dominant plasmid type in a (biased) dataset. On the other, careful experiments that characterise how one specific plasmid behaves are used to extrapolate beyond their scope. Both approaches need to be weighed in the balance to understand plasmid biology. Notably, the leading region as a concept arose from experimental studies of the F plasmid but is now commonly defined bioinformatically, as we have done here. Our finding that CAI is higher in the first ~25kb of the transfer strand applies to the F plasmid and many other Enterobacterales plasmids, but not all: plasmids in PTU-C and PTU-HI1A (both within MOB-H) lacked any clear signal in CAI. Canonical anti-defense genes like *psiB* have been noted to be present only in narrow-range plasmids, functioning in a species-specific manner (53). Existing knowledge of anti-defense genes is highly imperfect, but it is notable that PTU-C has very few known anti-defense systems in the leading region (most cluster at 50-100 ORFs) and PTU-HI1A had no complete anti-defense systems detected at all; this suggests alternative anti-defense mechanisms, but without knowledge of these it remains unclear how a plasmid like R27 (first isolated from *S. enterica*) is able to conjugate with high frequency into other Enterobacterales (54). Plasmids can lack the conjugation machinery but still be mobilizable, but 94% of plasmids with MOB-H relaxases are conjugative, in stark contrast to MOB-F and MOB-P where mobilizable plasmids are significantly more prevalent and persistent (55). Potentially the apparent deficit of anti-defense genes in MOB-H is connected to this difference.

Our findings emphasise the need to analyse plasmids in cohesive units such as PTUs. It is important to appreciate the differences in genome organization and evolutionary strategy between these units. The leading region of the F plasmid is GC-rich and its genes have high CAI. So too do those of many – but not all – plasmids, with clear differences by relaxase type. This demonstrates that some canonical features of a leading region do not apply to all conjugative plasmids. Although the *oriT* of PTU-C plasmids has been experimentally validated, it has been suggested that they may carry a second *oriT* (56). The lack of a peak in CAI in its apparent leading region makes this seem possible. If so, the F plasmid might be an unreliable guide to the general dynamics of conjugative transfer. Recent pioneering work on the F plasmid has shown that the temporal dynamics of the leading region are complex and exquisitely controlled (10,42,43). It seems likely that other plasmids may have similarly sophisticated but potentially different adaptations; it remains important to characterise plasmids beyond the well-studied minority.

## Acknowledgements

This article originated from a group project during the EMBO Workshop ‘Plasmids as Vehicles of AMR Spread’ held in Trieste, Italy 18-22 September 2023. We would like to thank Enas Newire and Adrian Cazares Lopez for their involvement in the original group project. We are grateful to the workshop organisers and other participants for discussions and feedback on our early ideas and results on RM systems. We would also like to thank three anonymous reviewers for their comments on a previous version of this paper.

## Data availability and methods

A github repository with analysis is available at https://github.com/liampshaw/Leading-region-adaptation. Associated data is available via Zenodo (doi: 10.5281/zenodo.20040901).

## Supplementary material

**Table S1**. Metadata for plasmids analysed in this study, including the subset of plasmids where we were confident in the location of the leading region (n=1,751).

**Table S2**. Detection of RM systems with known targets after screening random genomes from the 37 genera in our initial study.

**Table S3**. Anti-defense and defense hits detected in plasmids with DefenseFinder against HMM profiles (evalue<10^−10^).

**Table S4**. Codon weights for species analysed in this study, calculated from the codon usage of highly expressed genes in reference genomes.

**Table S5**. Codon adaptation index for all plasmid genes (using optimal *E. coli* weights) as well as their functional annotation with eggnog-mapper.

**Figure S1.**
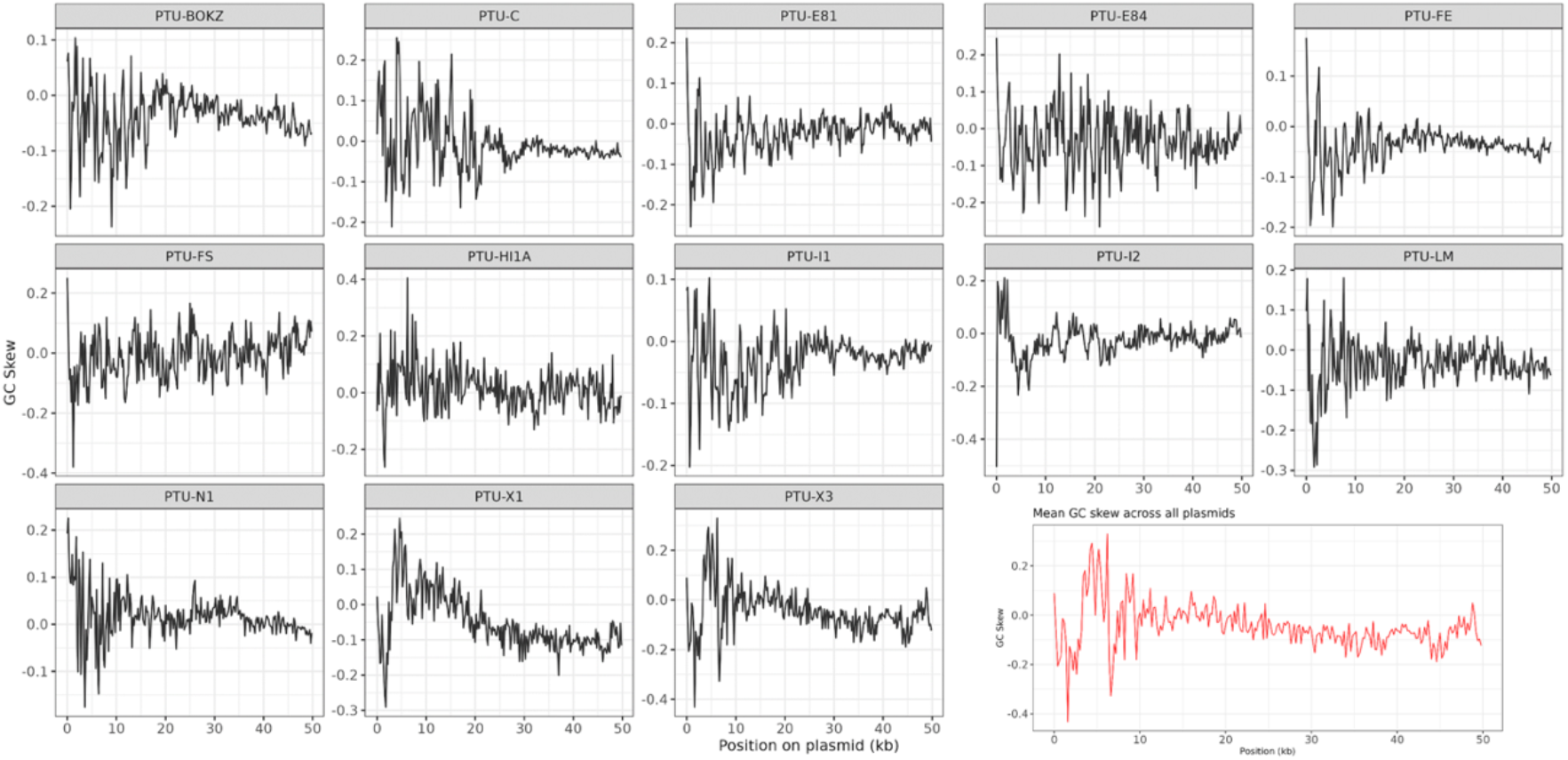
GC skew in the first 50kb of different PTUs. Sliding window of 200bp across 13 PTUs (1,751 plasmids), with all plasmids orientated so their leading region starts at zero.

**Figure S2.**
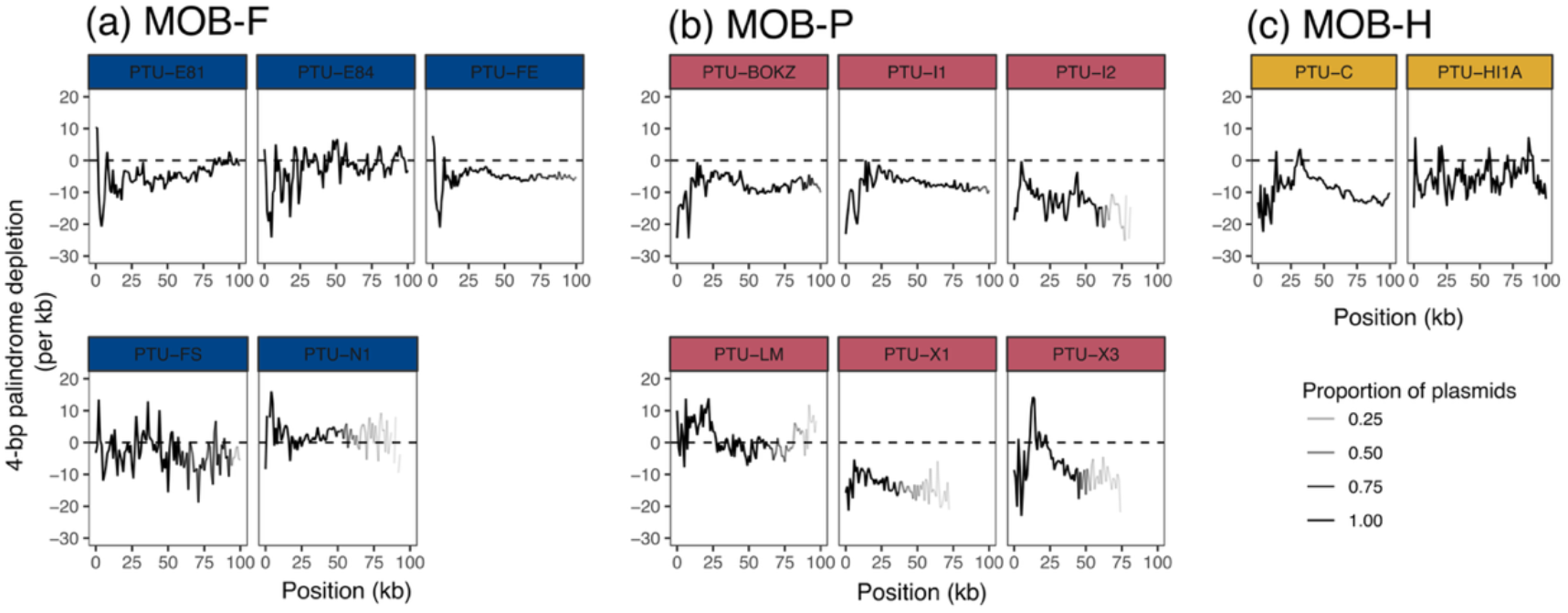
4-bp palindrome density across plasmids.

**Figure S3.**
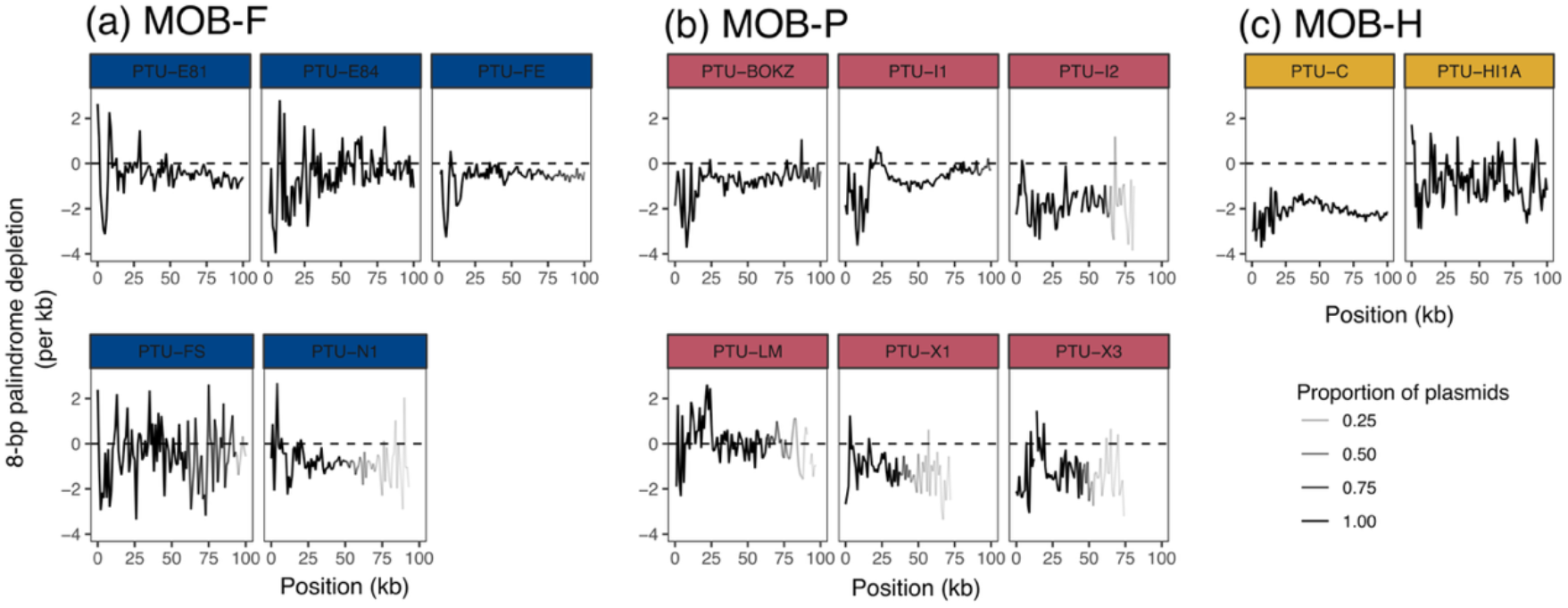
8-bp palindrome density across plasmids.

**Figure S4.**
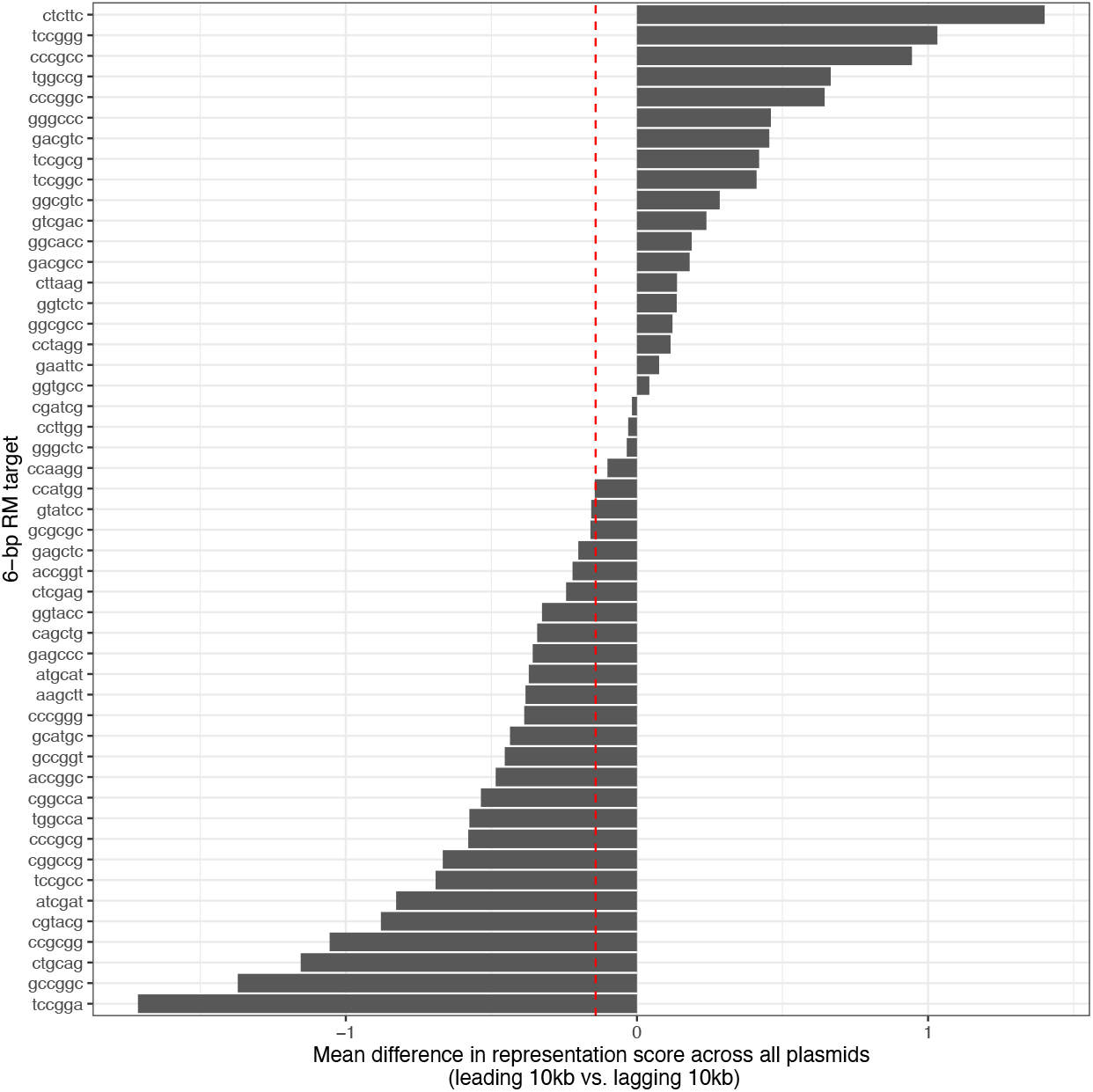
Depletion of 6-bp RM targets in leading vs. lagging region, averaged across all plasmids. Bars represent the mean difference in RMES score between the leading and lagging regions across all plasmids (10kb each), with negative scores corresponding to more evidence of depletion in the leading region after correcting for sequence composition. The red dashed line is the mean of the means (−0.14; Wilcox signed rank test p=0.083).

**Figure S5.**
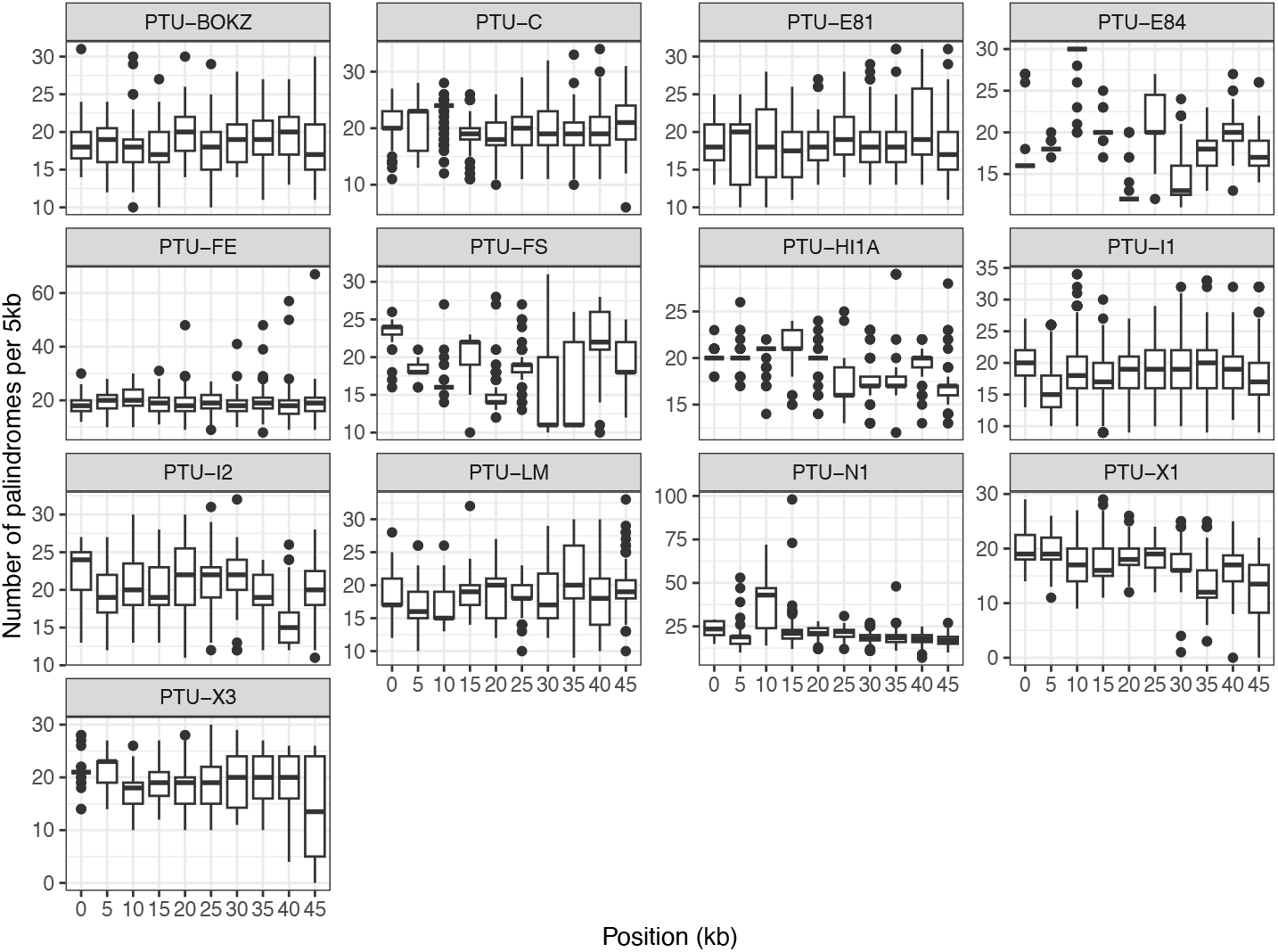
No variation in the number of near-perfect palindromes >4bp with plasmid position. Results are shown binned by 5kb for the first 50kb of each PTU.

**Figure S6.**
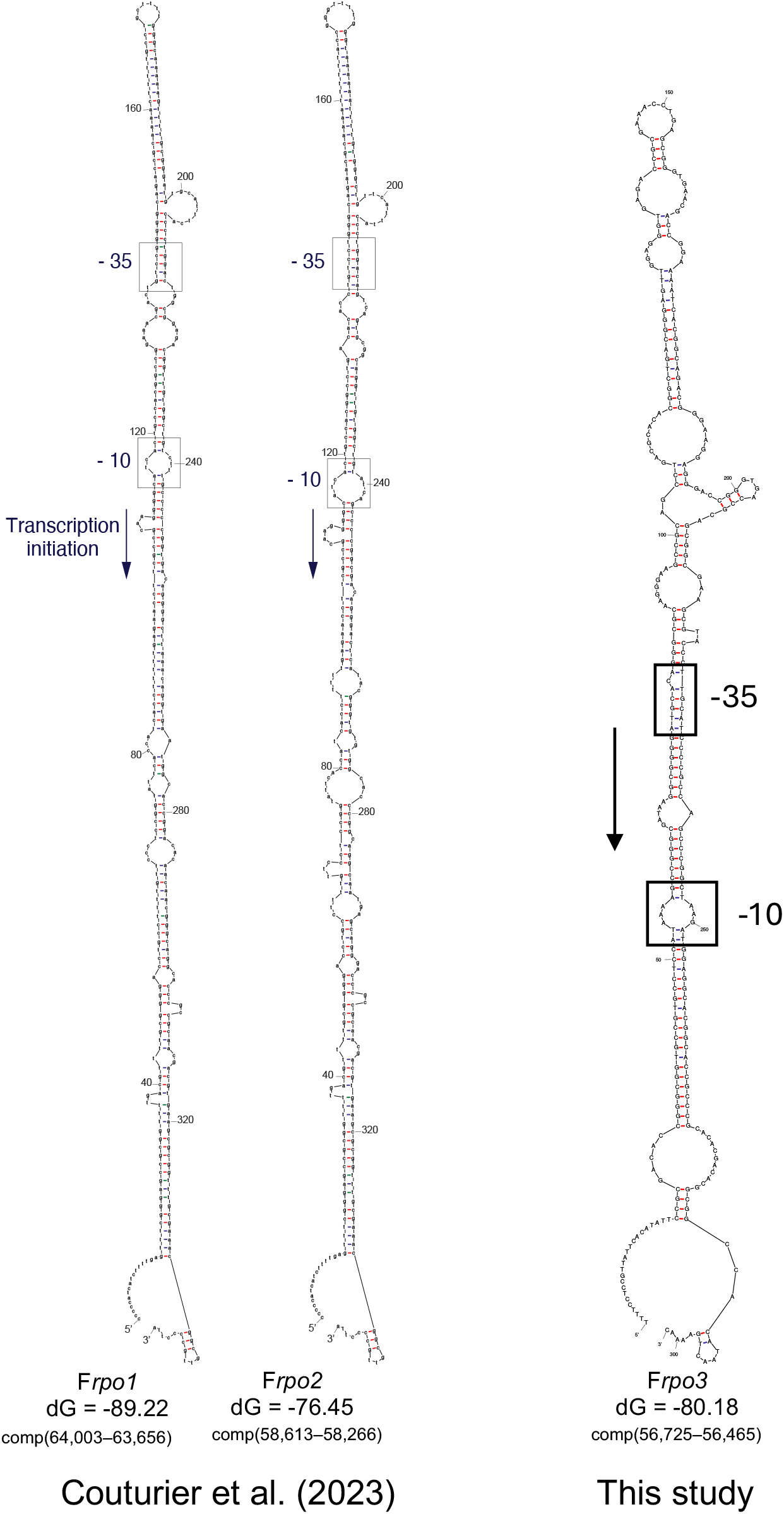
Frpo3 is a single-stranded DNA promoter. Secondary structures predicted with mFold v3.6 for three single-stranded DNA promoters in the F plasmid. Coordinates are with respect to reference sequence NC_002483.1, ‘comp’ denotes reverse complement. This region was independently identified as a ssDNA promoter by Wen et al. (42) and its function explored further by Fraiken et al. (43).

